# Single Cell Transcriptome Conservation in Cryopreserved Cells and Tissues

**DOI:** 10.1101/067884

**Authors:** Amy Guillaumet-Adkins, Gustavo Rodríguez-Esteban, Elisabetta Mereu, Alberto Villanueva, August Vidal, Marta Gut, Ivo Gut, Holger Heyn

## Abstract

A variety of single cell RNA preparation procedures have been described. So far these protocols require fresh starting material, hindering complex study designs. We describe a sample preservation method that maintains transcripts in viable single cells and so allows to disconnect time and place of sampling from subsequent processing steps. To demonstrate the potential, we sequenced single cell transcriptomes from >1,000 fresh and conserved cells. Our results confirmed that the conservation process did not alter transcriptional profiles. This substantially broadens the scope of applications in single cell transcriptomics and could lead to a paradigm shift in future study designs.

## Background

Within complex tissues cells differ in the way their genomes are active. Despite the identical DNA sequence of single cells, their distinct interpretation of the genetic sequence makes them unique and defines their phenotype [1]. While in many complex biological systems cell type heterogeneity has been extensively analyzed in molecular and functional experiments, its extent could only be estimated due to the technical limitation to assess the full spectrum of variability. With the advent of single cell genomics, cell type composition can be deconvoluted for unprecedented insights into the complexity of multicellular systems. Exemplarily, single cell transcriptomics studies resolved the neuronal heterogeneity of the retina [2], the cortex and the hippocampus [3,4], but also advanced our definition of hematopoietic cell states [5]. Moreover, single cell genomics studies shed light on cellular relationships in dynamic processes, such as embryo development [6] and stem cell differentiation [7]. Here, it was the assessment of hundreds to thousands of single cell gene expression signatures that allowed the determination of tissue composition at ultra-high resolution. In addition to providing insights into the complexity of the analyzed samples, single cell studies provide an invaluable resource of biomarkers that define cell types [3,8] or differentiation states [9].

Different single cell RNA sequencing techniques allow the quantification of minute transcript amounts from up to thousands of single cells, however, their exclusive dependence on fresh starting material strongly affected study designs [10]. In particular, the need for immediate sample processing hindered complex study setups, such as time course studies, or sampling at locations without access to single cell separation devices. Indeed, seminal work on the composition of complex systems was performed with readily accessible tissues from model organisms and the extent to which conclusions can be projected to human physiology is unclear [2,3,5].

Here we evaluate a sample preservation method that disconnects time and location of sampling from subsequent single cell processing steps. It enables complex experimental designs and widens the spectrum of accessible specimens. Specifically, samples were cryopreserved to maintain cellular structures and the integrity of RNA molecules for single cell separation months after archiving. To demonstrate the potential of our method, we analyzed single cell transcriptomes from 1,418 fresh or cryopreserved cells, obtained from cell lines or primary tissues.

## Results and Discussion

Cell integrity and RNA quality present crucial requirements for successful single cell transcriptome sequencing experiments. Conventional conservation processes, such as freezing, lead to crystallization and disruption of cellular membranes, which impedes subsequent single cell preparation. To conserve intact and viable cells for cell and tissue archiving, cryoprotectants are commonly used, however, their applicability for single cell experiments has not been established. We tested weather cells preserved with the cryoprotectant dimethyl-sulfoxide (DMSO) are suitable for single cell genomics workflows. To this end, we sequenced 675 fresh and 743 cryopreserved single cells derived from cell lines and primary tissues prepared with the MARS-Seq protocol [5,11] (**Fig. S1**). Subsequently, a variety of statistical methods, including the most common measures in single cell genomics, were applied to determine potential systematic biases introduced by the conservation method.

**Fig. 1.**
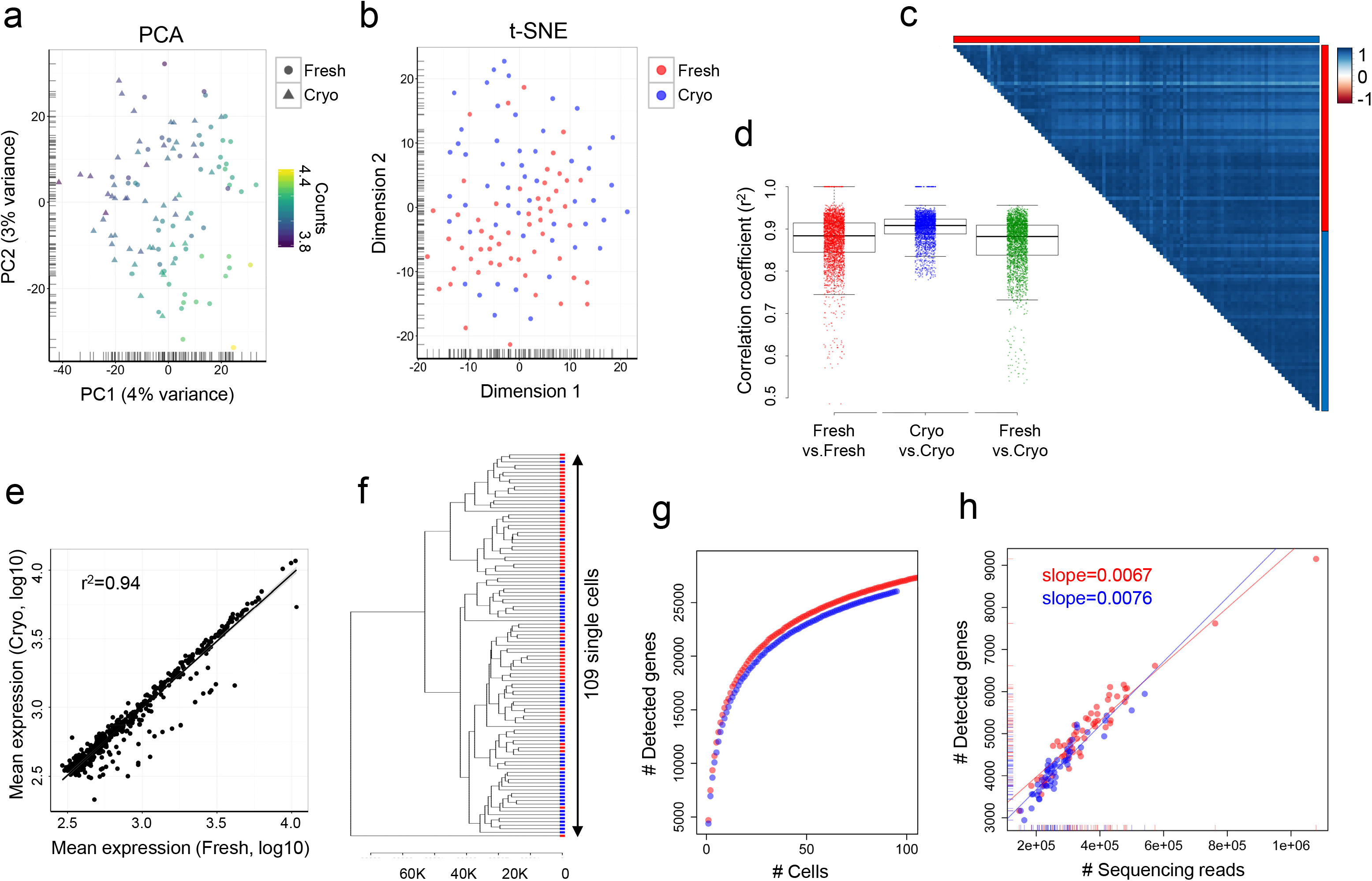
Comparative analyses of single cell transcriptome data from fresh and cryopreserved HEK293 cells. (**a**) Gene expression variances between fresh (circles) and cryopreserved (triangles) cells are displayed as principal component analysis (PCA) of scaled log2-transformed transcript counts indicating the sequencing depth (log10 of total counts). (**b**) A t-distributed stochastic neighbor embedding (t-SNE) representation of fresh (red) and cryopreserved (blue) cells and scaled log2-transformed transcript counts. (**c**) Pearson’s correlation analysis between fresh (red) and cryopreserved (blue) cells displaying the correlation coefficient (r^2^). Single cells were ordered by their position on the 384-well plate. (**d**) Distribution of Pearson’s correlation coefficients (r^2^) within and between processing conditions. The median coefficients are indicated. (**e**) Linear regression model comparing average gene expression levels of the 500 most expressed genes in fresh and cryopreserved cells. The coefficient of determination (r^2^) is indicated. (**f**) Unsupervised hierarchical clustering of single cells based on the 500 most expressed genes. (**g**) Cumulative gene counts split by fresh (red) and cryopreserved (blue) cells and analyzed using randomly sampled cells (average of 100 permutations). (**h**) Comparative analysis of the number of sequencing reads and detected genes using a linear model. The slope of the regression line was calculated separately for fresh (red) and cryopreserved (blue) cells.

To determine potential impacts of the cryopreservation procedure on single cell RNA profiles, we isolated single cells from four cell lines HEK293 (human embryonic kidney cells), K562 (human leukemia cells), NIH3T3 (mouse embryo fibroblasts) and MDCK (canine adult kidney cells) by fluorescence-activated cell sorting (FACS). The cells were either freshly harvested or cryopreserved at -80°C prior to the single cell separation. To minimize technically introduced batch effects both conditions were processed simultaneously for library preparations and sequencing reactions. We sequenced our single cell transcriptomes significantly deeper than previous MARS-Seq studies [5,11], which allowed us to detect biases introduced by the cell preparation method (**Fig. S1**).

Satisfyingly, the transcriptional profiles of conserved cell line samples were indistinguishable from freshly processed cells in commonly used dimensionality reduction representations (**Fig. 1a,b** and **Fig. S2**). Principal component analyses (PCA) and t-distributed stochastic neighbor embedding (t-SNE) showed a homogeneously mixed population of single cells, a result also observed with independent data normalization techniques (**Fig. S2** and **S3**). We found sequencing depth to be the main source of variance between single cells, a common confounding factor in single cell transcriptome studies (**Fig. 1a**) [12]. High similarities between single cells of both conditions were confirmed by correlation analysis showing highly consistent and representative gene expression profiles after cell conservation (**Fig. 1c-e** and **Fig. S4**). As expected analyzing homogenous cell populations, expression profiles showed high correlation values between single cells of the same type and condition (Pearson’s correlation test, median r^2^:0.88-0.91). However, also between conditions transcription profiles were highly correlated (Pearson’s correlation test, median r^2^:0.88), suggesting the freezing process to conserve single cell transcriptome profiles (**Fig. 1c,d**). Consistent expression profiles were further supported by highly correlating mean expression values when directly comparing both conditions (**Fig. 1e**). Also, hierarchical clustering of single cells using the 500 most variable genes could not distinguish between conditions, further suggesting equal gene expression profiles between fresh and conserved cells (**Fig. 1f** and **Fig. S4**). In order to evaluate potential impacts on comparative expression analyses involving fresh and conserved sample types, we assessed differentially expressed genes between both conditions (**Fig. S5**). We could not detect any significantly differentially expressed transcripts between fresh and cryopreserved sample (adjusted p-value > 0.05), supporting the possibility to include conserved material in studies profiling freshly processed samples.

Following the exclusion of systematic biases introduced by the conservation method, we evaluated to which extent biological information is maintained within single cells. We compared gene information and associated biological processes between both conditions. Here, a comparable number of genes was detected by accumulating information from single cells, suggesting that the power to detect gene transcripts in the conserved material is not reduced (**Fig. 1g** and **Fig. S6**). Indeed, we noticed that mainly sequencing depth influenced the total number of detected genes per experiment. The conservation process did not show a loss of gene expression information. In line, we found a similar linear relationship between the number of sequencing reads and detected genes for both conditions (linear regression model, slope: 0.0067 and 0.0076; Fig. 1h and Fig. S6). This provides further support for the capacity to extract the same transcript information at a given sequencing depth.

Biological processes that one might suspect to change due to a challenge, such as cell cycle and apoptotic programs remained unchanged (Fig. 2a,b). Moreover, both conditions were suitable for the identification of biologically relevant information. We detected comparable numbers of cells with active cell cycle programs in fresh and conserved cells, identified by functional single cell clustering [13] and the identification of indicative processes (e.g. sister chromatin segregation) and marker genes (e.g *CCNB1*) (**Fig. 2c,d**).

**Fig. 2.**
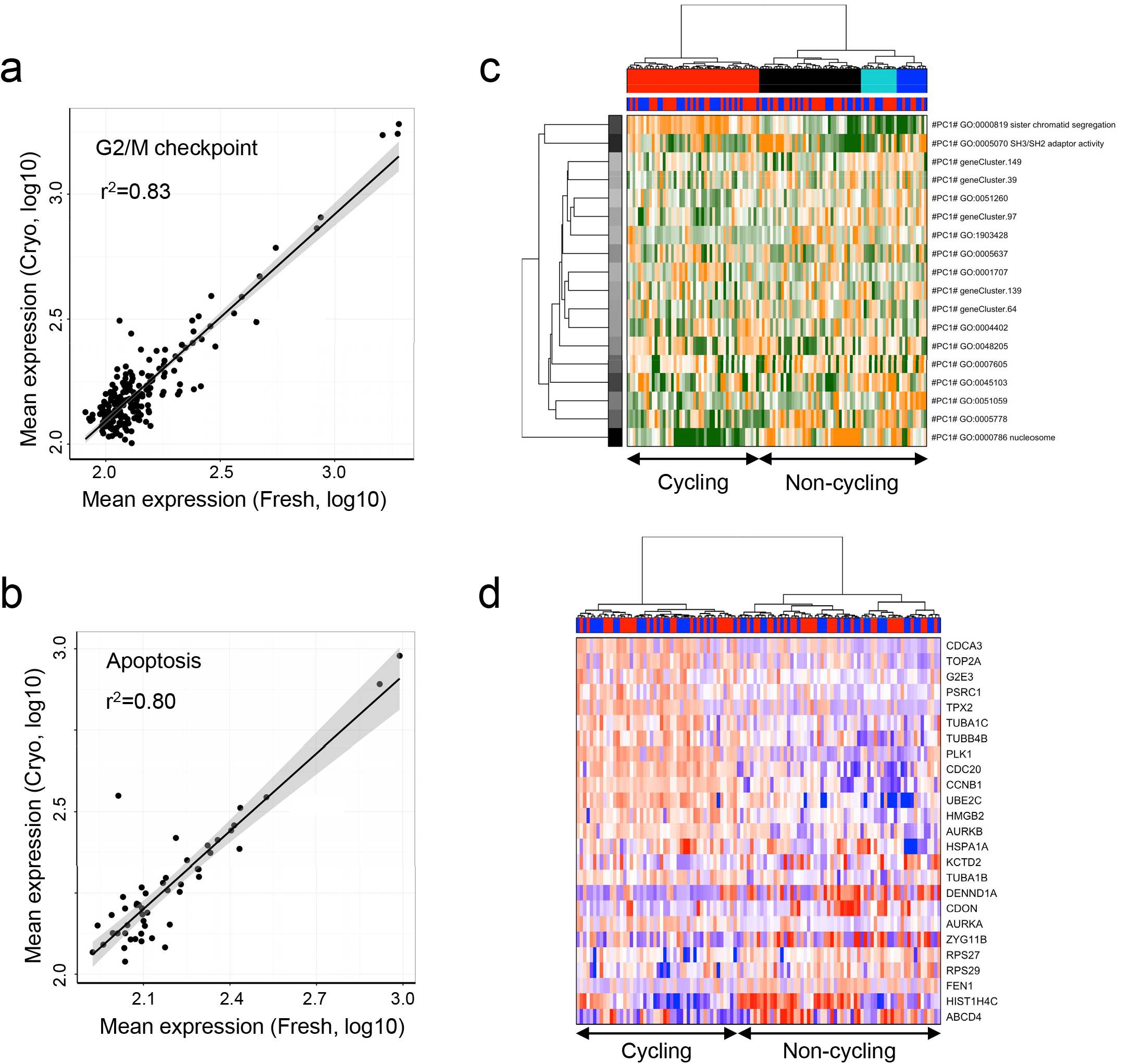
Cryopreservation does not affect functional analysis of HEK293 cells. (**a,b**) Linear regression analysis for average expression levels of (**a**) G2/M checkpoint or (**b**) apoptosis related genes [23] comparing fresh and cryopreserved cells. The coefficient of determination (r^2^) is indicated. (**c**) Hierarchical clustering of fresh (red) or cryopreserved (blue) single cells based on transcriptional programs (defined by gene ontology) and correlating genes (referred to as gene clusters) [13]. Displayed are the most variable aspects (rows), their importance (row colors) and the most enriched annotation (row labels). The cell cycle status determined by functional annotation and marker genes is indicated for the two dominant cell clusters. Transcriptional programs and gene clusters are summarized (orange: overrepresented; green: underrepresented). (**d**) Hierarchical clustering of fresh (red) and cryopreserved (blue) single cells based on transcriptional programs and correlating gene sets (as in c). Displayed are the 25 most informative genes and the cell cycle status is indicated for the two dominant cell clusters. Gene expression levels are displayed as relative intensities following transformation [13] (red: high; blue: low).

Although conserving cell cultures for single cell analysis opens up the applicability to more complex experimental designs, we intended to further widen the application spectrum to complex primary tissues. We performed MARS-Seq experiments on freshly resected and cryopreserved mouse colon tissues and subsequently extended the work to human tumor samples. A fresh mouse colon sample was split. One part was cryopreserved for one week before single cell separation. No significant differences between fresh and conserved colon cells following transcriptome sequencing were found, consistent with results from the cell line experiments (**Fig. 3a-d**). Gene information was unaltered (**Fig. 3e,f**) and sufficient to derive biologically relevant information. We were able to identify transit amplifying (TA) cells, secretory enteroendocrine cells and differentiated enterocytes in both conditions (**Fig. 3g**), major cell types present in the colon mucosa. The single cell transcriptome data enabled us to clearly assign colon cell types to cell clusters using marker genes [8], such as *Reg4* (secretory cells), *Apoa1* (enterocytes) or ribosomal proteins (TA cells) (**Fig. 3h**). We conclude that the conservation process did not alter the transcriptional profile of single cells and that both, single cell sequencing of fresh and conserved tissues is equally suitable to extract biologically relevant information, such as cell type specific programs.

**Fig. 3.**
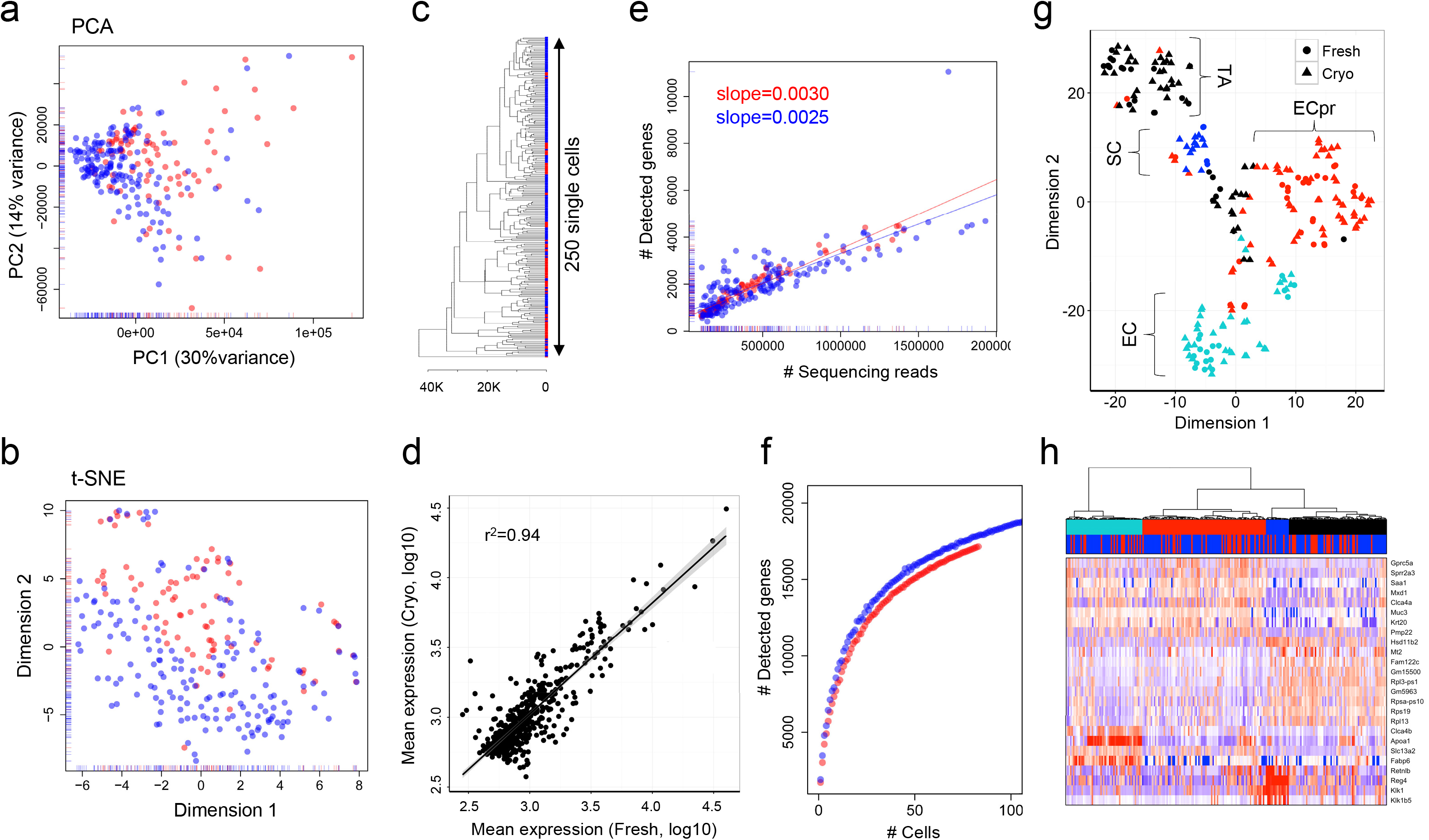
Comparative analyses of single cell transcriptome data from fresh (red) and cryopreserved (blue) mouse colon cells. Gene expression variances between cells are displayed as (**a**) PCA or (**b**) t-SNE representation of normalized transcript counts. (**c**) Unsupervised hierarchical clustering of single cells based on the 500 most expressed genes. (**d**) Linear regression analysis comparing average gene expression levels of the 500 most expressed genes of fresh and cryopreserved cells. The coefficient of determination (r^2^) is indicated. (**e**) Comparative analysis of the number of sequencing reads and detected genes using a linear model. The slope of the regression line was calculated separately for fresh (red) and cryopreserved (blue) cells. (**f**) Cumulative gene counts split by fresh (red) and cryopreserved (blue) cells and analyzed using randomly sampled cells (average of 100 permutations). (**g**) t-SNE representation derived from transcript counts of functionally selected genes [13]. Fresh (circles) and cryopreserved (triangles) conditions are indicated. Cell types were annotated based on marker gene set (derived from Grün et al. [8]) expression analysis (TA: transit amplifying, SC: secretory cells, EC: enterocytes, ECpr: enterocytes precursors). (**h**) Hierarchical clustering of fresh (red) or cryopreserved (blue) single cells based on transcriptional programs (defined by gene ontology) and correlating genes (defined as gene clusters). Displayed are the 25 most informative genes and gene expression levels are displayed as relative intensities (red: high; blue: low) [13].

Finally, we successfully applied our method to a patient-derived orthotopic xenograft (PDOX) cryopreserved for three months before processing. A freshly resected ovarian clear cell carcinoma orthoxenograft was processed simultaneously with the matched cryopreserved PDOX. Consistent with prior observation, single transcriptome profiles of fresh or conserved tumor cells did not differ in their transcriptional profiles (**Fig. 4a-d**), further highlighting tissue conservation to be possible for various experimental designs, including tumor samples.

**Fig. 4.**
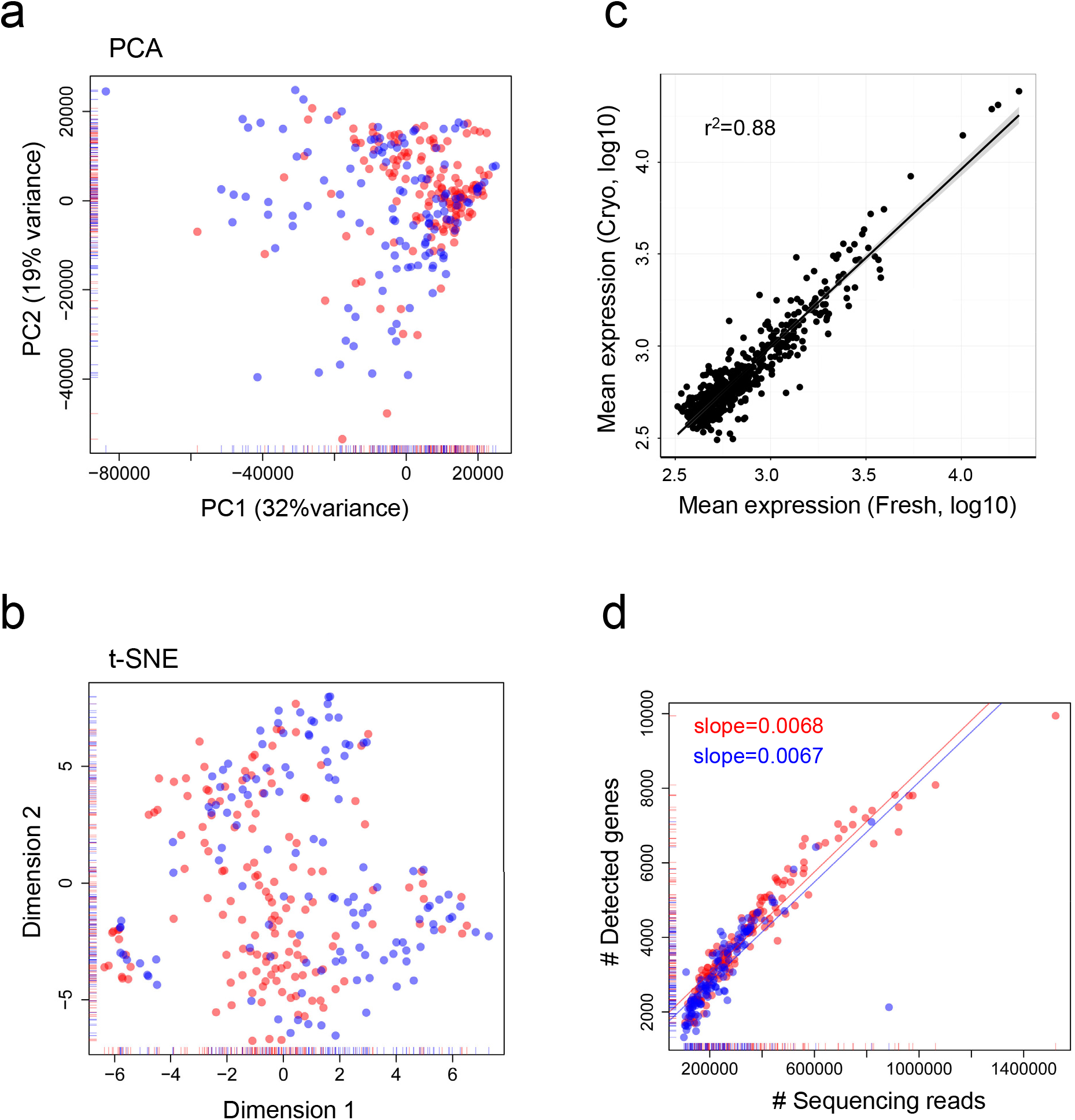
Fig. 4 Comparative analyses of single cell transcriptome data from fresh (red) and cryopreserved (blue) orthotopic ovarian tumor xenograft cells. Gene expression variances between cells are displayed as (**a**) PCA or (**b**) t-SNE representation of normalized transcript counts. (**c**) Linear regression analysis comparing average gene expression levels of the 500 most expressed genes of fresh and cryopreserved cells. The coefficient of determination (r^2^) is indicated. (**d**) Comparative analysis of the number of sequencing reads and detected genes using a linear model. The slope of the regression line was calculated separately for fresh (red) and cryopreserved (blue) cells.

## Conclusions

Using the here established cryopreservation method single cell transcriptome profiles from cells and tissues did not differ from freshly processed material. The method constitutes a straightforward and powerful tool to broaden the scope of single cell genomics study designs. Importantly, cryopreservation can be readily implemented into standard single cell genomics workflows, without modifications in established protocols. In addition, although combined here with the MARS-Seq technique, other single cell RNA-Seq methods are likely to result in similar outcomes [14,15]. Although recent work described the value of nuclear RNA analysis [4,16], the content from viable cells is likely to result in a more complex transcriptome description for in depth study.

Taking into account that cellular structures are conserved during cryopreservation, different downstream applications, including genome or epigenome sequencing might also benefit from this method. Cryopreservation was previously described to conserve open chromatin structures in ATAC sequencing experiments [17], pointing to a wide application spectrum of cryopreserved material.

In conclusion, the conservation process we present here does not modify transcriptional profiles of single cells taken from cell culture or tissues. Cells cryopreserved by our method are equally well suited as fresh cells to extract relevant biological information, such as cell type specific programs. This substantially broadens the scope of applications in single cell transcriptomics and could constitute a paradigm shift for single cell study designs.

## Methods

### Cell Line Sample Preparation

Human cell lines HEK293 (human embryonic kidney cells) and K562 (human leukemia cells) were acquired from the German Collection of Microorganisms and Cell Cultures (DSMZ). NIH3T3 (mouse embryo fibroblasts) and MDCK (canine adult kidney cells) were kindly provided by Dr. Manel Esteller (IDIBELL, Spain). HEK293, NIH3T3 and MDCK were maintained in DMEM (10% fetal bovine serum, FBS; 1% Penicillin/Streptomycin) at 37°C (5% CO2). K562 suspension cells were cultured in RPMI (10% FBS; 1% Penicillin/Streptomycin) at 37°C (5% CO2). For cryopreservation, cells were trypsinized, pelleted and resuspended in freezing solution I (10% DMSO; 10% heat-inactivated FBS; 80% DMEM) or solution II (10% DMSO; 90% non-inactivated FBS). Subsequently, cells were frozen with gradually decreasing temperatures (1°C/min) to -80°C (cryopreserved). For single cell analysis, cryopreserved cells were rapidly thawed in a water bath with continuous agitation and placed into 25 ml of cold 1x HBSS. Fresh cells were trypsinized, pelleted and resuspended in 1x HBSS. Before sorting, cells from both conditions were filtered (70 μm nylon mesh) and propidium iodide staining identified dead/damaged cells. To avoid batch effects fresh and cryopreserved single cells were sorted into the same plates and distributed over both sequencing pools.

### Primary sample preparation

Female athymic nu/nu mice (Harlan) between 4 to 6 weeks of age were housed in individually ventilated cages on a 12 hour light-dark cycle at 21-23°C and 40-60% humidity. Mice were allowed free access to an irradiated diet and sterilized water. Primary mouse colon was dissected from an athymic nu/nu mouse and placed on ice. The sample was divided and half of the colon was immediately prepared for single cell separation, while the other half was minced on ice, placed into freezing solution II (10% DMSO, 90% non-inactivated FBS) and frozen with gradually decreasing temperatures (1°C/min) to -80°C (cryopreserved). After storage for one week at -80°C, the sample was rapidly thawed in a water bath in continuous agitation and placed into 25 ml of cold 1x HBSS. For single cell separation the fresh and conserved samples were minced on ice and enzymatically digested in 5 ml 1x HBSS and 83 μl collagenase IV (10,000 U/ml) for 10 min at 37°C. Single cells were separated by passing the sample through a 0.9 mm needle and filtration (70 μm nylon mesh). Cells were washed once in ice cold 1x HBSS and resuspended in DMEM before sorting. Dead and damaged cells were identified by propidium iodide staining. For practical reasons (tissue derived from one single mouse), fresh and cryopreserved single cells could not be sorted into the same plate.

### Orthotopic ovarian carcinoma engraftment

To analyze matched fresh and cryopreserved viable tumor samples, we generated an ovarian orthotopic tumor model, referred as Orthoxenograft® or patient-derived orthotopic xenograft (PDOX). Therefore, we implanted a primary clear cell ovarian carcinoma into the ovaries of athymic nu/nu mice (matched organ of origin). Briefly, the primary tumor specimen was obtained at the Hospital Universitari de Bellvitge (Barcelona, Spain). The selected patient had not received cisplatin-based chemotherapy. Non-necrotic tissue pieces (~2-3 mm^3^) from a resected clear cell ovarian carcinoma were selected and placed into DMEM, supplemented with 10% FBS and 1% penicillin/streptomycin at room temperature. Under isofluorane-induced anesthesia, animals were subjected to a lateral laparotomy, their ovaries exposed and tumor pieces anchored to the ovary surface with prolene 7.0 sutures [18,19]. Tumor growth was monitored 2 to 3 times per week. When the tumor grew, it was harvested and cut into small fragments. Subsequently, it was transplanted into a new animal and cryopreserved at -80°C as viable tumor (as described above). After 107 days, the tumor was newly resected from the mouse and processed together with the matched cryopreserved sample (maintained at -80°C) for single cell separation and sorting. The morphology of the primary tumor and the engrafted tumor was compared by H&E staining in paraffin-embedded sections. For single cell separation the sample was rapidly thawed in a water bath in continuous agitation and placed into 25 ml of cold 1x HBSS. For single cell separation the fresh and conserved samples were minced on ice and enzymatically digested in 5 ml 1x HBSS and 83 μl collagenase IV (10,000 U/ml) for 15 min at 37°C. Single cells were separated by passing the sample through a 0.9 mm needle and filtration (70 μm nylon mesh). Cells were washed once in ice cold 1x HBSS and resuspended in DMEM before sorting. In order to enrich human cells during the sorting procedure, tumor cells were stained for 1 hour at 4°C with α-EpCam (CD326, eBioscience, 1:100). Propidium iodide staining identified dead/damaged cells. To avoid batch effects fresh and cryopreserved single cells were sorted into the same plates and distributed over both sequencing pools.

### Library preparation and sequencing

To construct single cell libraries from polyA-tailed RNA, we applied massively parallel single-cell RNA sequencing (MARS-Seq) [5,11]. Briefly, single cells were FACS-sorted into 384-well plates, containing lysis buffer and reverse-transcription (RT) primers. The RT primers contained the single cell barcodes and unique molecular identifiers (UMIs) for subsequent de-multiplexing and correction for amplification biases, respectively. Spike-in artificial transcripts (ERCC) were added at a dilution of 1:16x10^6^ per cell. PolyA-containing RNA was converted into cDNA as previously described and then pooled using an automated pipeline (liquid handling robotics). Subsequently, samples were linearly amplified by *in vitro* transcription, fragmented, and 3’-ends were converted into sequencing libraries. The libraries consisted of 192 single cell pools. Multiplexed pools (2-6) were run in one Illumina HiSeq 2500 Rapid two lane flow cell following the manufacturer’s protocol. Primary data analysis was carried out with the standard Illumina pipeline. We produced 52 nt of transcript sequence reads for the cell lines and the mouse colon tissue and 83 nt for the tumor xenograft sample.

### Data processing

The MARS-Seq technique takes advantage of two-level indexing that allows the multiplexed sequencing of 192 cells per pool and multiple pools per sequencing lane. Sequencing was carried out as paired-end reads; wherein the first read contains the transcript sequence and the second read the cell barcode and UMIs. Quality check of the generated reads was performed with FastQC quality control suite. Samples that reach the quality standards were then processed to deconvolute the reads to single cell level by de-multiplexing according to the cell and pool barcodes. Reads were filtered to remove polyT sequences. Sequencing reads from human, mouse or canine cells were mapped with the RNA pipeline of the GEMTools 1.7.0 suite [20] on the genome references for human (Gencode release 24, assembly GRCh38.p5), mouse (Gencode release M8, assembly GRCm38.p4) and dog (Ensembl v84, assembly CanFam3.1). We adapted the GEMTools RNASeq pipeline for its usage on single cell reads. The analysis of spike-in control RNA allowed us to discard reads from empty wells or damaged cells. Cells with less than 10^5^ reads or more than 2x10^6^ reads were discarded. Gene quantification was performed using UMI corrected transcript information to correct for amplification biases.

### Data analysis

To estimate systematic biases introduced by the conservation technique, single cells from both conditions were compared using two commonly used data pre-processing strategies and different metrics to assess similarities between cells. Statistical analyses shown in this manuscript were carried out using R, version 3.3.0. Functions referred below belong to the R stats package when not indicated otherwise.

#### Transformation and scaling

Fresh and cryopreserved data sets were filtered for low-quality cells (<800-1,800 genes) and genes (<10 cells and <20 transcript counts). The overlapping genes of both data sets were merged, resulting in a joint gene-cell matrix for each experiment. Principal component analyses (PCA) and t-distributed stochastic neighbor embedding (t-SNE) were performed using log2-transformed counts-per-million (with a prior count of 1 as values to use for expression) and the *scater* package. Both methods classify in an unsupervised manner by grouping most similar cells into clusters, however, the t-SNE algorithm also captures non-linear relationships. To determine differentially expressed genes, cell measurements have been modeled as a mixture of negative binomial and Poisson distributions, using the *scde* package [21]. As batch effects are a source of variation, variability introduced during the experimental phase (e.g. sequencing pools) has been taken into account. The Bayesian approach behind the method allows gene expression inferences from amplified and drop-out events. To fit cell models we used the default implementation. The quality of the models was evaluated with the value of correlation with the expected magnitude, which was positive for all cells. Further, the distribution of drop-out events for each sample appeared highly and negatively correlated to the expression magnitude, showing the value 1 associated to zero magnitudes. The differential expression analysis of the cell line datasets revealed that the relative contribution of each gene between the two groups of cells was highly comparable (adjusted p-value>0.05). Differences between gene expression profiles were studied by correlating relative or absolute gene counts and by displaying the most expressed genes. In linear regression models, we found a strong linear correlation (r^2^ ~ 0.9) between the means of the two groups (considering the number of non-zero cells to compute mean values) and a strong agreement of relative count levels for the 50 most expressed genes (Fig. S5).

#### Library size normalization

With the aim to preserve the original data structure, gene expression based on UMIs was scaled to correct for differences in library sizes between cells. Library size normalization, although a very simple normalization method, has shown to perform better for single cell RNAseq data than more sophisticated normalization approaches developed for bulk RNAseq and comparable to single cell specific normalization algorithms [22]. PCA on the gene expression matrix was performed using the *prcomp* function. The t-SNE clustering was performed on the gene expression matrix with the library *Rtsne* and “perplexity” parameter was set to the total number of cells divided by five. For subsequent analyses, following filter steps were applied: genes represented by ≤1 UMI in a given cell and genes present in ≤50% of the cells were discarded. Pearson’s correlation matrices were calculated with the *cor* function and represented using the *corrplot* library (**Fig. 1c**). Hierarchical clustering was performed with the *hclust* function with complete linkage on a euclidean distance matrix, calculated with the *dist* function for the 500 most expressed genes. Clustering was represented as dendrogram with the *dendextend* library 1.2.0.

### Subpopulation and heterogeneity analysis

In order to functionally interpret transcriptional heterogeneity between single cells, expression profiles of HEK293 and mouse colon cells were modeled using Pathway and geneset overdispersion analysis (PAGODA) [13], included in the *scde* package. Models were constructed similar to the differential expression analysis and the goodness of fits has been successfully assessed for all cells. A correction for pool compositions and sequencing depth was performed. PAGODA allows the identification of principal aspects of heterogeneity, capturing the most overdispersed gene sets and normalizing for undesired aspects. We applied Gene Ontology, *de novo* and custom pathways to define clusters of gene sets (aspects). Subsequently, cells were clustered based on a weighted correlation of genes driving these aspects [13]. The most variable aspects or genes were displayed in the hierarchical clustering highlighting the most significant genes, GO terms or *de novo* gene sets. Further, distances from the hierarchical clustering were used to visualize cells in two dimensions through a t-SNE plot. In both hierarchical clustering and t-SNE plots, we could not detect evidence for differences between fresh and cryopreserved samples. To assign cell types to single cell clusters, we used colon specific custom gene sets and marker genes (e.g. *Apoa1, Reg4* or ribosomal genes) derived from Grün et al.[8]. Cell cycles states were defined by marker genes (e.g. *CCNB1)* and Gene Ontology term enrichment (e.g. sister chromatid segregation). Apoptosis (Hallmark_Apoptosis; M5902) and G2/M (Hallmark_G2/M_CHECKPOINT; M5901) gene sets were derived from the GSEA database [23].

## List of abbreviations

(DMSO): Dimethyl-sulfoxide

(ERCC): External RNA control consortium

(FBS): Fetal bovine serum

(FACS): Fluorescence-activated cell sorting

(GSEA): Gene set enrichment analysis

(MARS-Seq): Massively parallel single-cell RNA sequencing

(PAGODA): Pathway and geneset overdispersion analysis

(PDOX): Patient derived orthotopic xenograft

(PCA): Principal component analyses

(t-SNE): T-distributed stochastic neighbor embedding

(UMIs): Unique molecular identifiers

## Ethics

All animal protocols were reviewed and approved according to regional Institutional Animal Care and Use Committees. The primary tumor specimen was obtained at the Hospital Universitari de Bellvitge (Barcelona, Spain). The study was approved by the Institutional Review Board and written informed consent was collected from the patient.

## Author’s contribution

HH conceived and directed the study. AGA implemented and applied sequencing library preparation protocols. GER established single cell RNA sequencing processing pipelines and performed statistical analysis with significant support of EM. AlV and AuV led the PDOX tumor models and contributed primary mouse and human samples. HH wrote the manuscript with support from MG and IG. All authors read and approved the final manuscript.

## Acknowledgements

We thank the cytometry unit of the CCiT (University of Barcelona) for successfully implementing single cell sorting.

